# Morphometric reconstructions atlas shows insult-driven plasticity in cortical VIP/ChAT interneurons

**DOI:** 10.1101/2020.08.25.263178

**Authors:** Nadav Yayon, Oren Amsalem, Amir Dudai, Or Yakov, Gil Adam, Marc Tessier-Lavigne, Nicolas Renier, Idan Segev, Michael London, Hermona Soreq

**Affiliations:** The Edmond and Lily Safra Center for Brain Sciences (ELSC), The Life Sciences Institute, The Hebrew University of Jerusalem, Jerusalem, 9190401, Israel; The Departments of Biological Chemistry, The Life Sciences Institute, The Hebrew University of Jerusalem, Jerusalem, 9190401, Israel; The Departments of Neurobiology, The Life Sciences Institute, The Hebrew University of Jerusalem, Jerusalem, 9190401, Israel; Depratment of Biology, Bass Biology Building, Stanford, CA 94305, USA; Sorbonne Université, Paris Brain Institute, INSERM, CNRS, 75013, Paris, France

## Abstract

We developed an automatic morphometric reconstruction pipeline, Pop-Rec, and used it to study the morphologies of cortical cholinergic VIP/ChAT interneurons (VChIs). Cholinergic networks control high cognitive functions, but their local modulation and stress-driven plasticity patterns remained elusive. Reconstructing thousands of local VChIs registered to their exact coordinates in multiple cleared murine cortices highlighted distinct populations of bipolar and multipolar VChIs which differed in their dendritic spatial organization. Following mild unilateral whisker deprivation, Pop-Rec found both ipsi-and contra-lateral VChI dendritic arborization changes. Furthermore, RNA-seq of FACS-sorted VChIs showed differentially expressed dendritic, synapse and axon-modulating transcripts in whisker-deprived mice. Indicating novel steady-state morphological roles, those genes also clustered distinctly in naïve single cell VChIs. This VChIs “morpheome” atlas is the first example of unbiased analysis of neuronal populations and holds the possibility to compare neuronal structure-function relationships across experimental conditions.

## Introduction

Neuronal morphology defines electrophysiological properties and the spatial distribution of synapses. Therefore, the morphology of dendritic and axonal arbors is a major determinant of neuronal specialization and integration in circuits ^1,2^. Morphology is among the major determinants used to categorize neuron types, and efforts to accurately reconstruct a neuron’s complete shape dates back to the beginning of modern neuroscience^3–5^. As a result, continuous efforts have been devoted over time to precisely define the most salient morphological parameters (e.g. dendrite diameter and length, the number of neurites per cell and axonal span)^3,6–8^. Neuro-morphology features are notably subject to dynamic changes, for example in response to stressful experiences^9,10^, viral infection^11^ or disease states^12^. However, our understanding of the stability of neuronal branches is hampered by the difficulty to obtain large-scale population level statistics of morphological changes following biological and experimental perturbations. This is primarily due to the time-consuming manual/semi-manual nature of the task and the need for very high-resolution imaging to precisely reconstruct neurons. Compared to other high-throughput techniques (e.g. Single-Cell/Nuclei RNA-Seq^13,14^), manual neuron reconstructions offer limited statistical power and fail to detect fine cumulative differences in neuronal features across experimental conditions. Furthermore, limited numbers of reconstructions per mouse hamper the prospects of correcting for sample variability and batch effects in neuronal population studies. Recent improvements in tissue clearing^15–17^, light-sheet imaging^18^ and analysis techniques^19–22^ now streamline the acquisition of entire neuronal populations over large intact brain regions. However, performing automatic reconstructions from cleared tissues is still complicated because of two major issues: 1) inhomogeneities in labeling, signal-to-noise ratios and tissue background, which can introduce variance and statistical noise in the reconstructions; and 2) the limited resolution of light sheet microscopes, which prevents the reconstruction of most neuron types due to their high density. To address these problems, we developed a novel tool, Pop-rec, which we applied to the large-scale reconstruction of a tractable neuronal population, the cortical cholinergic neurons.

The mammalian cortex receives cholinergic input from two major sources: deep nerve growth factor (NGF)-responding basal nuclei projection neurons^23^ and local cortical interneurons expressing vasoactive-intestinal polypeptide (VIP) and choline acetyl-transferase (ChAT)^24^: the VIP/ChAT interneurons, or VChIs. Relatively little is known about the morphometry of VChIs, which have been described as sparse, bipolar neurons mostly distributed in cortical layer 2/3^25,26^. Single neuron RNA-sequencing (RNA-seq) analyses have indicated the reactiveness of VChIs to immune-related neurokines^27^. Notably, VChIs were shown to co-release GABA and acetylcholine(ACh), and recent findings highlight their diverse capacities to modulate the activity of neighboring neruons^28,29^. Recently, we demonstrated the functional involvement of VChIs in modulating cortical somato-sensory responses *in-vivo* and showed that they receive direct input from nucleus basalis projection neurons^26^ and react to inflammatory signals^27^. However, the structural diversity of these VChIs and its long-term plasticity remain largely unknown.

To investigate the morphological diversity of VChI neurons in control and whisker-deprived mice, we used Pop-Rec, an analysis pipeline that enables unbiased automatic and quantitative reconstructions. We applied Pop-Rec to the study of VChIs in the mouse barrel cortex receiving the primary inputs from the whiskers^28^. Notably, the barrel cortex responses are modulated by cholinergic signaling^30,31^ and whole brain profiling demonstrated that it is especially enriched with VChIs^25^. This feature enabled us to generate an atlas of VChI morphologies in the barrel cortex. We then tested if and how the morphology of these neurons is disturbed following mild, unilateral whisker deprivation. Finally, we established a list of regulated transcripts in VChIs barrel neurons following whisker deprivations, which outlined possible pathways maintaining the stability of their morphologies.

## Results

### The Pop-Rec pipeline

To specifically tag VChIs, we crossed C57/B6 mice expressing Cre recombinase under control of the endogenous ChAT gene (ChAT-IRES-Cre) with a tdTomato reporter line (Ai14, Jackson Laboratories) (Movie 1). We perfused littermates, isolated their cortices (Figure 1. a, b) and processed them according to the iDISCO+ protocol^19^ for imaging with a light sheet microscope and acquired both tissue autofluorescence and specific signals in the barrel cortex region.

**Figure 1.**
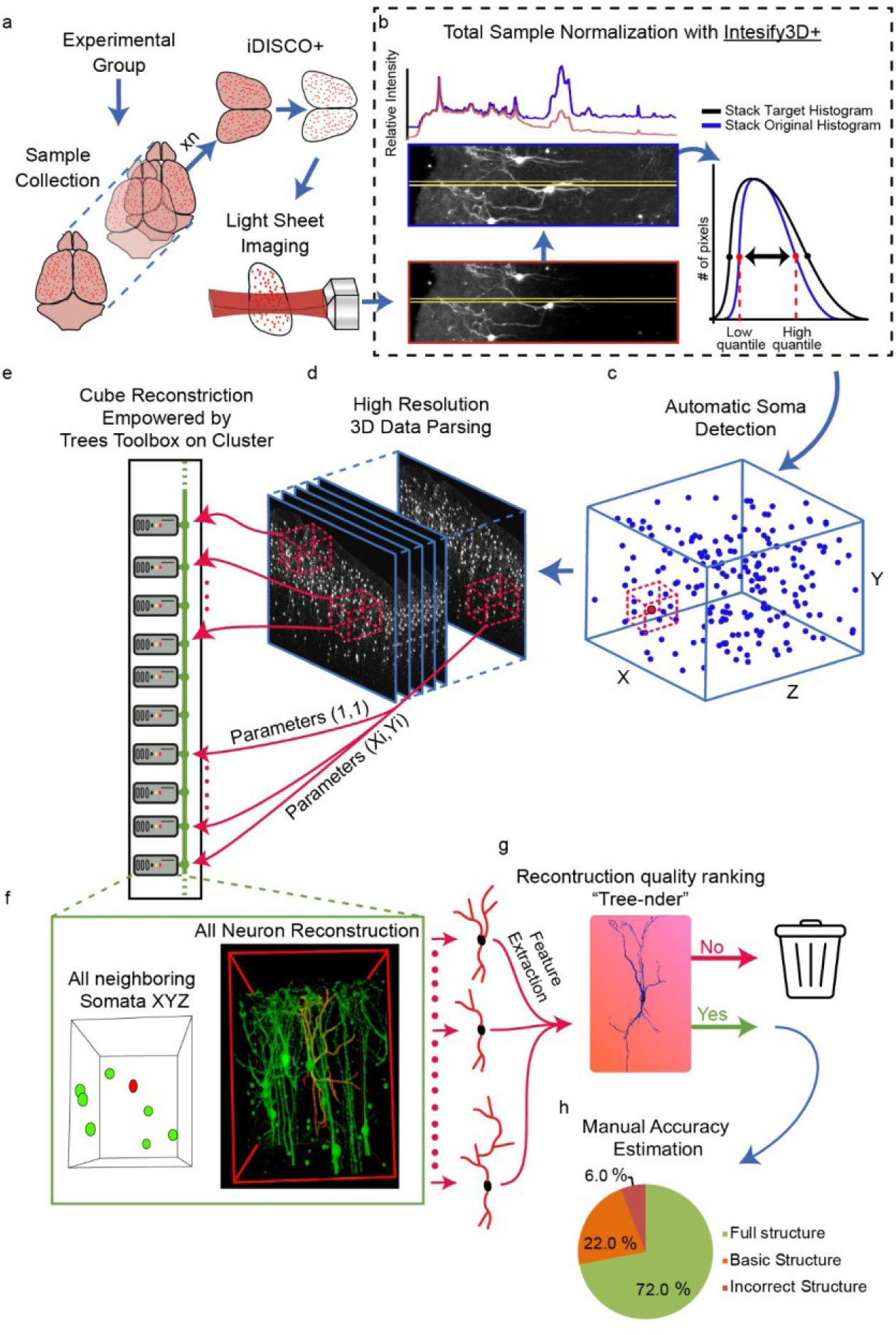
A scheme of the Pop-Rec analysis pipeline for automatic reconstructions. **a**. Chat-Cre x TdTomato mice were sacrificed and perfused, and cortices were isolated and subjected to iDISCO+ clearing and light-sheet imaging. **b**. signal intensity correction with intensify3D+. Depth intensity correction: Pre-red frame; Post-blue frame; Intensity profiles-yellow horizontal lines. Depth intensity correction followed by correction of full histogram equilibration across entire experimental biological sample set. **C** Automatic soma detection and coordinate tracking from the entire imaged volume. **d**. Data parsing into high-resolution mini-cubes around each cell together with neighboring cells and dendrites. **e**. Feeding every mini-cube to a computer cluster core together with a combination of reconstruction parameters (maximal gap distance and fluorescence threshold). **f**. Automatic reconstruction of center neuron (Architecture based on Trees Toolbox^20^ competitive reconstruction) while accounting for close proximity somata. **g**. Selecting an ideal parameter combination for each cell is done with a learning algorithm guided by NeuroM-defined features extracted from each version of the center neuron. Neurons that do not pass a certain quality threshold are discarded. **h**. Manual inspection of reconstruction accuracy (n=318).

Large scale tissue clearing and imaging introduces technical signal variability both within and between biological samples^32^. To compare variable dendritic signals of VChIs between cortical samples we constructed an updated version of our 3-dimensional normalization tool^32^ (Intensify3D+, Supplementary Figure 1) and corrected the observed signals within and between biological samples (Figure 1, c). Applying this algorithm to the imaged volume enabled automatic detection of VChI somata (Accuracy -- 97.94±0.012%, 14 cortical samples, Figure 1, c, Supplementary Table 2).

Large scale imaging datasets are challenging for current tracing and reconstruction tools which are often limited by imaging size and always require some form of manual intervention (BitPlane-IMARIS Filament Tracer neuroGPS^21^). To tackle this issue, we developed a data parsing approach by isolating “mini-cubes” that are large enough to contain the major dendrites of the center neuron as well as those of surrounding neurons in one mini-cube (Figure 1, c). To minimize large reconstruction errors, we fed a modified version of TreesToolbox^33^ information about the neighboring somata residing in the same cube as the center neuron. Together with somata location information, these mini-cubes are suitable for automatic reconstruction in a CPU cluster with a custom built input-output interface based on the Trees Toolbox^33^, as the reconstruction itself uses a heuristic competitive reconstruction approach which models the probability that colliding neurites belong to each soma. We further modified and adapted the Trees Toolbox to run in a cluster environment. Thus, all neurons could be simultaneously reconstructed, accounting for dendrites of neighboring neurons (Figure 1. f, g). From each mini-cube, we then discarded the surrounding reference neurons and kept only the center neuron. This process also maintained the original 3D coordinates of the reconstructed center neuron. Mimicking decision making of different human tracers and aiming to enhance the robustness of our reconstructions, we generated multiple reconstruction versions of each neuron with slightly different reconstruction parameters and with combinations of the maximal distance between two points and the fluorescent threshold in each version. In the last step of this reconstruction pipeline, we extracted multiple morphological features (NeuroM https://zenodo.org/record/209498#.Xl5xShMzbVs, Supplementary Table 1) from each version of the center neuron and ran an in-house regression-based learning algorithm which we called “Tree-nder” since it uses simple gestures (Right-Left) to quickly rank thousands of neuronal trees by the experimenter in a double blind, randomized manner across all samples in an experimental dataset (Figure 1h, Supplementary Figure 2). This step cleans out poorly reconstructed neurons, ranks each reconstruction version and automatically selects the best versions of high-quality reconstructions. It also excludes from further analyses neurons which do not cross the minimal quality threshold.

**Figure 2.**
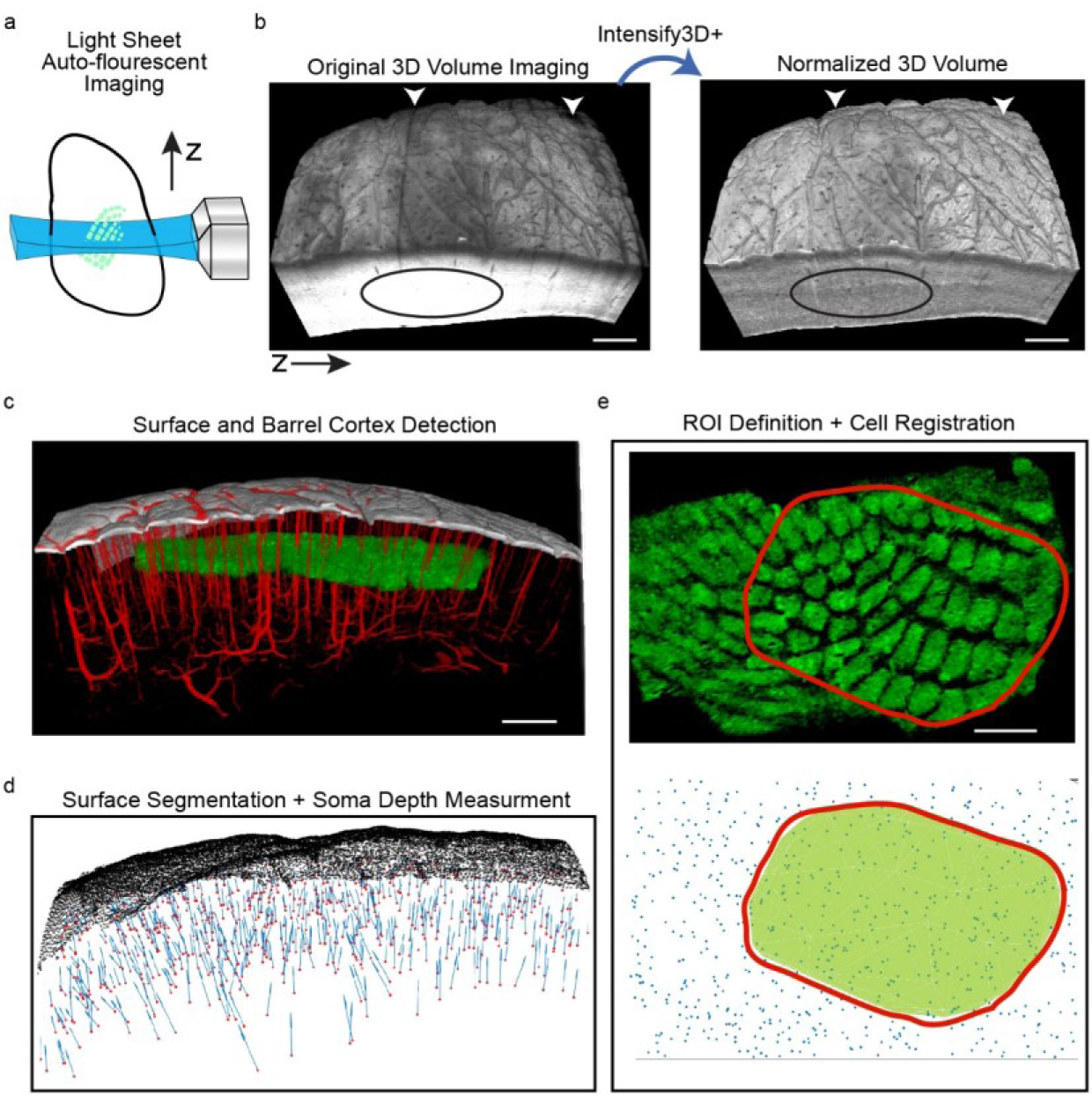
Auto fluorescence intensify3D+ normalization empowers Barrel cortex and surface detection. **a**. Scheme of light sheet auto-fluorescence imaging of barrel cortex region. **b**. Intensify3D+ normalization of pixel histograms corrects 3D light sheet scanning artifacts (white arrows) and intensity variability (black ellipse) in a tissue volume. **c**. Volume images post-normalization serve to isolate the cortical surface and estimate the boundaries of barrel cortex regions without any staining. Large blood vessels (red) could also be detected. **d**. 3D automatic detection and mesh segmentation of the cortical surface. The minimal Euclidean distance and direction (blue arrows) to the surface is calculated for each of the neuron somata (red dots). **e**. 3D barrel cortex ROI area (top – red line) is manually measured and segmented (FIJI^36^) to define an anatomical based ROI (bottom). This ROI is used to determine for each neuronal somata (blue dots) whether it is located within or outside of the barrel cortex.

To estimate the reconstruction accuracy and to evaluate a substantial fraction of the tested population we manually examined ∼300 out of ∼2400 neuron reconstructions from Chat-Cre X Ai14 mice. We found full length accuracy, meaning that the reconstruction reliably represented the major dendrites (proximal and distal arborizations) of 72% of these neurons and that we made no significant errors by including neighboring neuron details. In 22% of these neurons, the main dendritic structure was accurate, but some tracing errors occurred in distal dendrites, by assigning either too many or too few stem dendrites to the reconstructed neuron. Lastly, 6% of the neurons presented a significant missed tracing of a major stem dendrite (SD). Overall, the Pop-Rec approach hence yielded 94% accuracy in terms of basic neuron morphology and 72% accuracy in terms of the fine dendritic details (Figure h, Supplementary Table 2, Movie 2).

### Morphological classification of VChIs

The morphology and distribution of VChI neurons were recently described and categorized based on two distinct morphological sub-populations, bipolar/bi-tufted and multipolar^25,34,35^. We focused on inspecting the VChIs within the mouse barrel cortex, since they have been found to be enriched in somatosensory areas and are known to modulate sensory processing^25^. Recently, we have shown that despite their sparsity, VChIs can effectively modulate sensory processing in this cortical microcircuit^26^. To locate the barrel field, we normalized the auto-fluorescent signal from iDISCO+ cleared brains (Figure 2a) with Intensify3D+ and used this information to precisely measure the pial surface (top, dorsal) and the white matter tracts (bottom, ventral) boundaries of the cortex. Combining the above information with the locations of individual somata allowed us to register each neuronal soma inside the barrel cortex (Figure 2 b, c, Movie 3) and measure the cortical depth of these VchIs (Figure 2d).

After having established delineation of the barrel cortex and registration of VChIs within them, we next profiled the morphological diversity of VChIs within and between mice, by employing the Pop-Rec pipeline for 2383 VChIs in 14 cortical hemispheres from 7 mice (Figure 3a). Clustering of standardized morphological features (see Methods) by t-distributed stochastic neighbor embedding (tSNE) revealed two distinct morphological clusters, as indeed was suggested in previous reports^24,25^ (Figure 3b). Correspondingly, we identified two primary clusters, dominated by (i) cells with two principal stem dendrites (SD) termed – bipolar VchIs (biVChI) and (ii) cells with three or more principal stem dendrites termed multipolar VChIs (mVChI). Importantly, this segregation into two VChI subpopulations was consistent across individual mice, sexes and cortical sides (Supplementary Figure 3)

**Figure 3.**
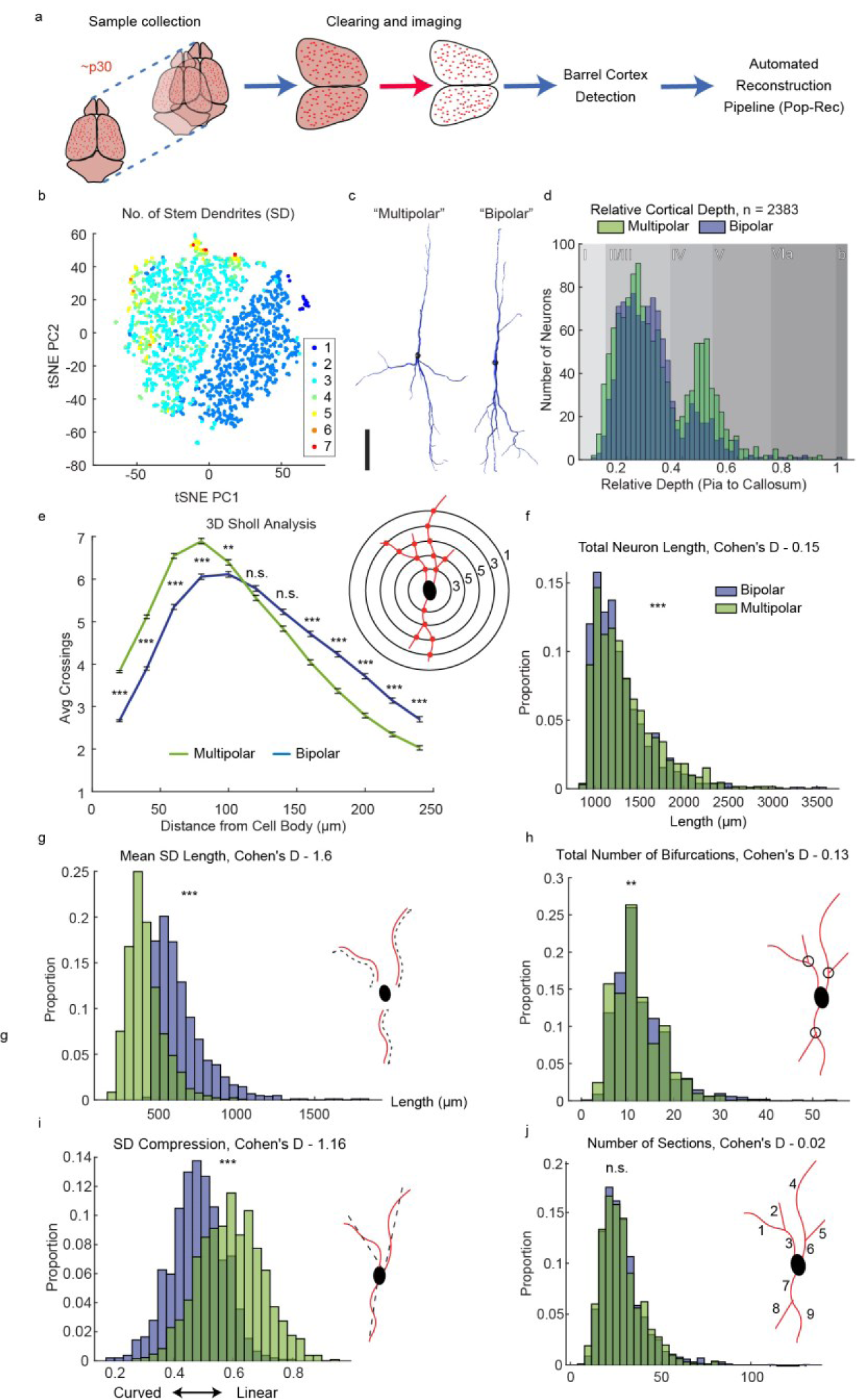
Discrimination of VChIs into bipolar and multipolar morphologies. **a**. Sample collection: naïve mouse cortices were processed (n = 7 mice, 14 cortex hemispheres), the barrel cortex area was defined, and VChIs were subjected to the Pop-Rec pipeline. **b**. tSNE clustering by morphological features. Colors represent numbers of stem dendrites or dendritic trees originating from the cell soma. **c**. Representative bipolar (left) and multipolar (right) VChIs of each group are shown. Horizontal scale bar - 100µm **d**. Cortical depth distribution of biVChIs (blue) MuVChIs (green) measured automatically by soma detection. **e**. 3D Sholl analysis of biVChIs and mVChIs (Bonferroni corrected linear mixed effects, LME). Example Sholl analysis and number of intersections (Right). Bars represent standard error of the mean. **f**. Distribution of biVChIs vs mVChIs for total neuron length. **g**. Distribution of biVChIs vs mVChIs for mean stem dendrite length per neuron. **h**. Mean number of bifurcations per neuron. **i**. Stem dendrite (SD) compression as a measure of dendrite curvature. **j**. Total number of sections per neuron. See Supplementary table 3 for full statistical description and LME models used in all panel unless stated otherwise. ***p<0.001, **p<0.01, *p<0.01. Cohen’s D < 0.2 = Small effect, 0.2<D<0.5 = Medium effect, D > 0.8 = Large effect.

Notably, biVChIs and mVChI differed fundamentally in various morphological features. First, the cortical distribution of these two subpopulations was distinct: biVChIs were homogeneously and almost exclusively distributed in layer 2/3, whereas mVChIs had a more significant proportion in layer 4/5. (Figure 3d). An independent 3D Sholl analysis^37^ of biVChIs and mVChIs, that was not included in the tSNE clustering showed less proximal and more distal branching in biVChIs, reaching up to 240µm from the cell soma. In comparison, mVChIs showed more proximal but less distal branching (Figure 3e). Accompanying the Sholl analysis, both the total length and stem dendrite (SD) compression were dramatically altered between the populations - the stem dendrites were shorter and more linear in mVChIs compared to biVChIs (Figure 3g, i). Strikingly, the overall total neuronal length, number of bifurcations and sections only slightly diverged between mVChIs and biVChIs (see Cohen’s D) (Figure 3f, h and j).

Together, these findings might indicate a metabolic constraint that limits the maximum average dendritic length, bifurcations and sections of VChIs at large. However, since these reconstructions are based on “mini-cubes” surrounding each neuron (500×350×350µm^3^), possible differences between these cell clusters could exist beyond 250µm from the soma. We conclude that barrel cortex biVChIs and mVChIs differ both in their cortical distribution as well as in their proximal and distal dendritic branching patterns, compatible with distinct innervation features and potentially unique functions of these two VChI types in the cortical microcircuit.

### VChIs in both contra- and ipsi-lateral cortices show morphological changes following sensory deprivation

In the first post-natal week of mouse pups, a loss of sensory inputs to the barrel cortex can trigger morphological changes to both barrel and septal neurons^10,38^. To characterize the effect of mild sensory deprivation on VChl neurons, we exposed young pups (p7) to a single event of light anesthesia with or without unilateral plucking of their full whisker pads. After 21 days we ran the PopRec pipeline (Figure 4a). Unlike in control mice, morphological clustering of experience-exposed mice failed to identify a clear segregation of VchI populations to biVChIs and mVChIs (Supplementary figure S4, See also Materials and methods). Seeking relevant differences in the structural features of this neuronal population within the barrel cortex, we pursued key features in the cortex of an un-deprived mouse, the ipsi-lateral cortex of a deprived mouse and the contra-lateral side of that mouse (full statistics in Supplementary table 4). We found a consistent morphological shift in several values: the SD compression, neurite section numbers and lengths, end and fork points and local tortuosity, all differed between the three groups (Figure 4b). To further test the sensitivity of our analysis at the level of the single neuron, we trained a linear regression model to predict the originating experimental group of each neuron, based on all of the morphological features (Supplementary table 4) (Figure 4c). This linear model succeeded in assigning the experimental group of single neurons with accuracy above chance and with a detection accuracy of 53% (chance at 37% as determined by mouse tagging permutations). Reflecting a whole brain impact, the best discernibility of this model emerged at the contralateral side of the deprived mice (Figure 4e).

**Figure 4.**
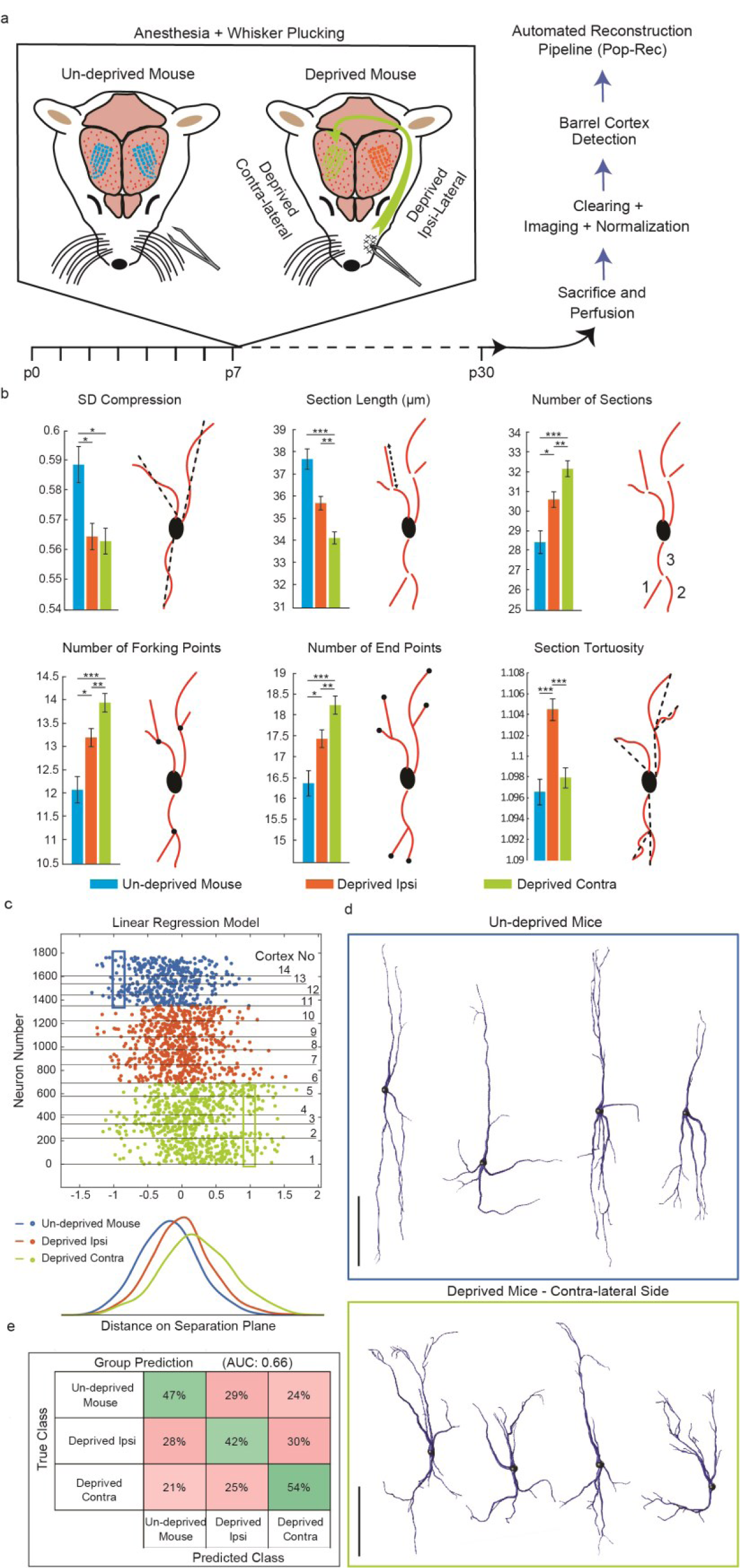
Effect of mild whisker deprivation on VChI morphology. **a**. Experimental scheme: mice were subjected to a single event of anesthesia with or without unilateral whisker plucking at p7. Mouse brains were collected at p30 and subjected to the Pop-Rec pipeline. **b**. Specific deprivation-induced changes in SD compression, section length and number of sections. Bars represent standard error of the mean. **c**. Linear regression model for separating single cells according to their morphological features. Neurons from distinct cortical samples are separated with horizontal lines. **d**. Top: representative VChI neurons from un-deprived (small blue rectangle) and bottom: deprived mice (small green rectangle). Scale bar: 100µm. **f**. Linear model performance summarized in a confusion matrix. Area Under the Curve: 0.66. See Supplementary table 4 for corresponding LME statistics. *P<0.05, **P<0.01, ***P<0.001. Data reflects 416, 658 and 692 analyzed VChIs from un-deprived mice, ipsi and contra-lateral cortices of deprived mice, respectively. Overall, 14 cortical samples (4 Un-deprived, 5 contralateral and 5 ipsilateral from 2 and 5 mice, respectively).

Overall, deprived VChIs had more dendrite sections with increased branching and less linearly oriented stem dendrites (Figure 4d). In summary, our neuro-morphology findings show that unilateral deprivation affects VChIs in both hemispheres, and that the difference between VChIs in deprived versus un-deprived mice is greater than between the ipsi- and contra-lateral hemispheres of deprived mice. Considering that the two cortical hemispheres are heavily connected by excitatory afferents^39^, this might reflect a brain-wide remodeling of connectivity.

### RNA-sequencing reveals modified dendrite structure-related transcripts following whisker deprivation

Our current understanding of the relationship between transcriptomic profiles and the morphological features of neurons is limited. Several genes are known to affect specific morphological features of axons, dendrites and dendritic spines, such as neurotrophic factors^40^ and transcription factors during development^41^. However, the transcriptomic response to whisker deprivation, and indeed the general links between specific neuronal structures and their transcriptomic profiles remained largely unknown. Since we observed specific morphological differences in dendritic structures in a mouse- and cortex-specific manner, we set out to determine if those differences reflected corresponding changes at the transcriptomic level. Given the sparsity of VChIs (which only constitute 0.5% of the neurons in the barrel cortex^26^) and their low survival rates following isolation using standard protocols, we developed a new isolation method based on a relatively short, light fixation followed by a Dounce-homogenization and digestion step similar, but not identical to that of single nuclei extraction procedures^13^. In contradistinction to the latter, our light fixation step enables preservation of fluorescently tagged cytoplasmic proteins and RNA, while sustaining a high yield and RNA quality of preserved neurons (Supplementary table 5). Performing this fixation approximately 2 minutes post-decapitation limits secondary transcriptomic responses (e.g. apoptosis and expression of immediate early genes) during the homogenization and isolation steps. Thanks to their intrinsic TdTomato labeling, VChIs isolated using this procedure could be detected and enriched using fluorescence-activated cell sorting (FACS). They further yielded high-quality RNA (RNA integrity number, RIN ∼8) via extraction with a dedicated kit with a calculated total RNA of ∼15pg RNA per neuron (Materials and Methods). Using this procedure, we isolated barrel cortex VChIs from both cortical hemispheres of mice that underwent unilateral whisker deprivation and controls (Materials and Methods). We then performed RNA library preparation and long RNA-sequencing using a dedicated low-input kit ((Figure 5a, b); Supplementary table 5, and Materials and Methods).

**Figure 5.**
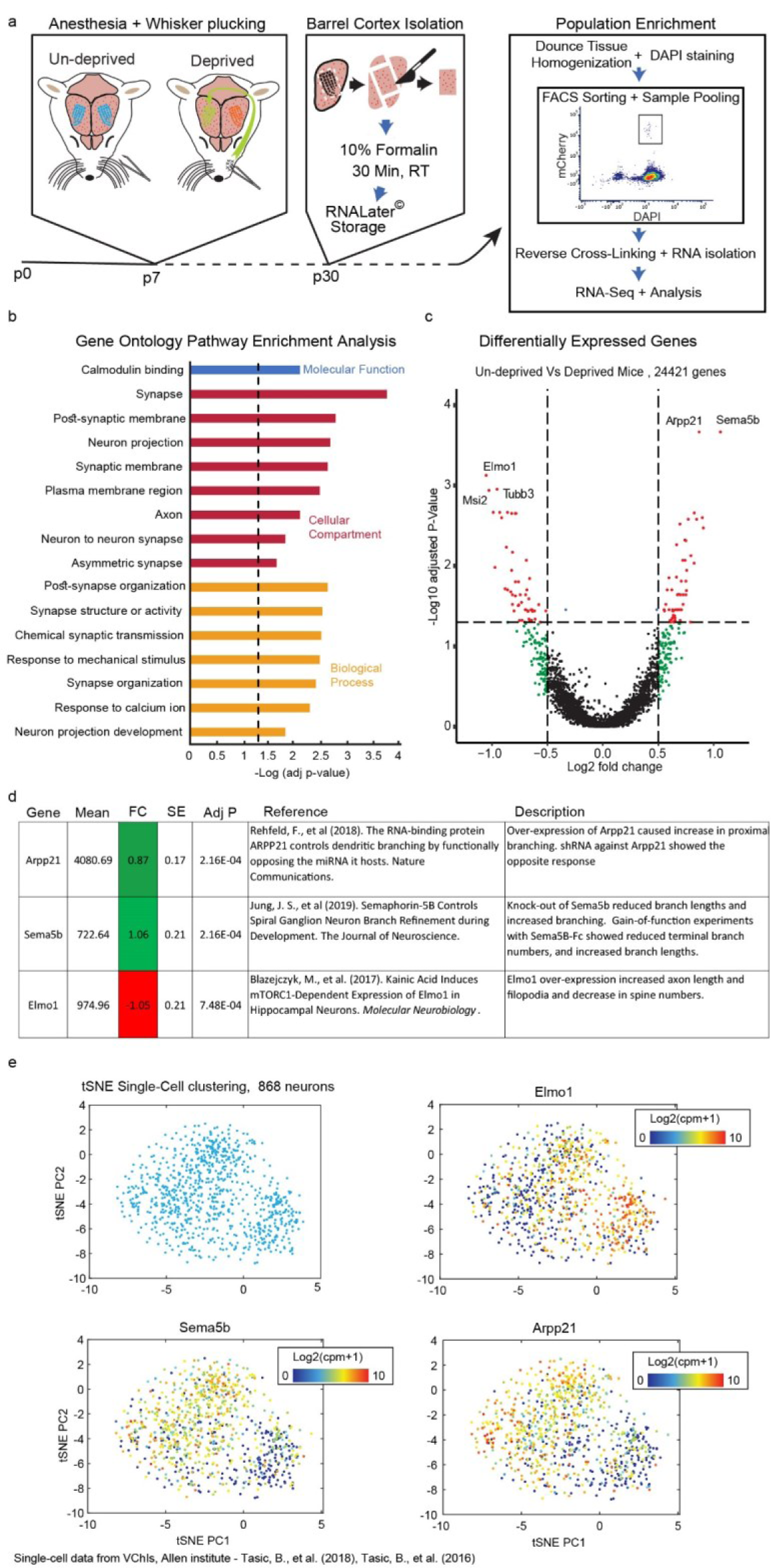
Transcript differences in enriched VChI populations following whisker deprivation. **a**. Experimental design of FACS-sorted RNA isolation strategy. Mice underwent whisker deprivation on day p7, were sacrificed on day 30 and their barrel cortices were isolated, lightly fixed, homogenized and subjected to fluorescence-based cell isolation (FACS). RNA was extracted from pooled -sorted TdTomato and DAPI-positive neurons from left and right cortices of different mice, subjected to RNA-Seq and the data analyzed. **b**. Gene ontology pathway enrichment analysis revealed specific cellular compartments and biological processes altered between control and deprived mice. Bar plots represent -log10 (Benjamini-Hochberg adjusted) p-values. **c**. Volcano plot depicting 110 differentially expressed transcripts between un-deprived (n = 4 pools, 8 samples – 4L, 4R) and deprived mice (n=6 pools, 12 samples – 6L, 6R). Positive change is more pronounced in deprived mice. **d**. Reported morphology-related function of Arpp21, Sema5b and Elmo1. **e**. tSNE clustering based on single nuclei RNA-seq data^42^ for expression profiles of Elmo1, Sema5b and Arpp21. Color scale: normalized log2(CPM+1) counts, n=868 single cell profiles.

Based on the concept that morphologically distict neurons present transcriptionally distinct expression patterns^13,43,44^, we predicted that RNA-seq profiles of specific neuronal populations in mice subjected to whisker deprivation might disclose those transcripts whose altered expression is reflected in the modified structure of those “experienced” neurons. We hence sought transcripts and pathways whose expression is altered 3 weeks post-deprivation; the time point at which we assessed the morphological changes using Pop-Rec (Figure 4a). Due to the sparsity of VChIs, we sequenced VChIs from several pooled cortices per sample (1-4 cortices). Interestingly, although the morphology of VChIs showed largely similar patterns between left and right cortices of untreated mice, we detected large differences between the transcript profiles of left and right cortices in untreated mice. By carefully controlling for the variability between transcriptomic profiles from pools and their distinct cortical sides (based on the Deseq2 model), we focused on identifying bilateral differences which were reproducibly detected as discriminating between deprived and control un-deprived mice.

Our extraction procedure enabled the detection an of over 14,000 trascripts per sample, with over 100 transcripts differentially expressed (DE) between deprived and un-deprived mice (Figure 5c). Supporting the neuronal source of these datasets, transcriptomics analysis confirmed the presence of numerous cholinergic (ChAT and VAChT) and VIP-related transcripts (Supplementary table 6). Gene ontology analysis of the DE transcripts reflected involvment of all major neuronal compartments, especialy ones related to synapse and neurite projections (Figure 5b) and of biological pathways related to synapse and neuron development and transcription factor networks. Intriguingly, the top two overexpressed genes, Arpp21 and Sema5b, and the top under-expressed gene, Elmo1, were very recently found to play substantial roles in the branching of dendrites and axons, as well as in synapse stability and microtubule organization^45–47^ (Figure 5d). In particular, over-expressed Arpp21 increases dendrite complexity and proximal branching^47^. Altogether, this analysis showed that mild whisker deprivation caused a shift in the transcriptional state of VChIs which might contribute to modifying their morphology and network communication capacities.

We were next interested in testing whether the changes in Arpp21, Sema5b and Elmo1 transcript levels might be reflected in the transcriptomic variability among VChIs in naïve mice. tSNE analysis of web-available single-cell VChI RNA-Seq datasets (Allen Institute) of 866 ACh-producing cortical neurons from naïve mice revealed segregation into at least two VChI transcriptomic clusters; importantly, this analysis highlighted the clear segregation of Arpp21, Sema5b and Elmo1 candidate transcripts into two distinct sub-clusters. Within these clusters, the Elmo1 transcript showed an inverse expression pattern to that of Sema5b and Arpp21, predicting divergent features similar to those observed for VChIs expressing these transcripts following whisker deprivation (Figure 5e). Taken together, these findings point to possible DE VChI transcripts that are functionally important for the transcriptomic-morphometric variability among VChIs in the murine brain.

## Discussion

In this study we constructed the structural “morpheome” of VChIs and combined neuro-morphology reconstructions of neuronal populations with RNA-seq transcriptomics from the same populations while accounting for experimental variability. We also sought to study the corresponding VChI differences between control and whisker-deprived mice, performed RNA-seq of the studied neuron populations and challenged our findings by mining web-available single cell transcriptomic datasets. Our findings enable exploration of the links between spatial resolution and intercellular differences at a single neuron resolution level, similar to transcriptomic spatial profiling^48,49^ and single-cell/nucleus RNA sequencing^20,50,51^. Adding high-throughput neuronal morphology analysis to these techniques will offer combined overview of the cellular proteome, transcriptome, genome, epigenome into our novel morpheome.

### Unbiased Population Reconstruction

Following the principles of single cell RNA-Seq, our structural analysis approach is based on the concept that the resolution of single neuron structures can be compromised to an extent, if sufficiently large cell numbers are available for comparative analysis. As this assumption only holds if the applied analysis is unbiased between all samples in the studied experimental group, we developed a computational correction approach for sample imaging bias (Intensify3D+) within and between biological samples. To enable rapid automated reconstruction of thousands of neurons with multiple structural parameters and handle huge imaging datasets, we “broke down” the data into small pieces that could then be analyzed in a cluster of CPUs. We defined the entire barrel cortex as our anatomically based region of interest (ROI) and subjected it to three-dimensional measurements of neuronal cortical depths and structures. As our approach keeps the original cortical coordinates of each neuron, it further enables regional analysis of groups of neurons such as those residing in specific cortical columns. Our strategy is amenable for use with many other, ever expanding automated reconstruction tools for single neurons, and our imaging (mini-cube) datasets can assist in benchmarking the performance of these algorithms in a well-controlled manner.

### The VChI morpheome in naïve mice

Previous studies have suggested the existence of two morphologically distinct VChI populations, proposed to reflect functionally different neuron subtypes. The relative proportion of bipolar vs multipolar VChIs is altered in different cortical regions^25^; also, higher numbers of multipolar VChIs are found in deep layers compared to bipolar cells^24^. In addition, the “Parent” VIP population has also been suggested to segregate according to stem dendrite number in the barrel cortex^52^. We show that unbiased clustering of tens of morphological features of reconstructed VChIs indeed reconfirms this dichotomy to bipolar and multipolar VChIs. In addition we also validated the previous observations that multipolar VChIs are relatively more abundant in deep cortical layers^24,25^. We identified different dendritic branching organization patterns and cortical distribution of bipolar (biVChIs) and multipolar (mVChIs) neurons. In contrast, we found only slight differences between the total dendritic length and total bifurcations of biVChIs and mVChIs, which might indicate that for a given metabolic state (RNA, Protein and lipid compositions), a neuron subtype could “decide” how to distribute its resources, but will always be limited in the maximum size and complexity of its extremities. Nevertheless, this altered economy might lead to distinct functional processing of the cortical microcircuit. To objectively pursue such a distinction while avoiding bias due to other objective measures (e.g. molecular, electro-physiological), one would have to sample and cluster the entire studied population and then prove by independent techniques that it indeed differs from others. We hope that our current atlas of morphometric VChIs will serve as a baseline for such future analyses, which may demonstrate if the terminal resources of neuronal extremities are a global neuronal feature. In addition, several axonal reconstruction of VChIs, indicate that their axons mostly occupy layers 4 and 5^24^. Further studies are needed to determine the detailed innervation patterns of bipolar and multipolar VChIs.

### Morphological response to whiskers deprivation

To detect small but cumulative changes in population morphologies, one must resolve sufficiently large numbers of dendritic resolution profiles from distinct structural states. We studied VChIs following a mild single event of whisker plucking (p7) which does not alter the barrel fields themselves^53^, and is often performed by mice as a social dominance signal and in the range of physiological manipulations^54^. Following whisker deprivation, VChIs showed more elaborate branching, with larger numbers but shorter lengths of dendrite sections. Interestingly, this feature was not altered in naïve mice between biVChIs and mVChIs. Future studies should investigate the relationship between branching and section length in terms of electrotonic properties of the neuron. Finally, we observed larger neuro-morphology differences between control and deprived mice than between the contra- and ipsi-lateral hemispheres of deprived mice. This may indicate that unilateral deprivation significantly affects the ipsi-lateral hemisphere as well, such that the impact of this and possibly other interference protocols on both the deprived and ipsi-lateral side and the entire cortex should be taken into consideration.

### VChI transcriptomic response to whisker deprivation

Robust enrichment of the sparse VChI populations from deprived and un-deprived mice by light fixation and FACS sorting enabled transcriptomic analysis, highlighting hundreds of differentially expressed genes and pathways in post-deprivation VChIs which might reflect altered VChI states. Of these, the top modified transcripts Arpp21, Sema5b and Elmo1 segregated into distinct clusters in single cell VChI profiles from naïve mice as well, showing parallel expression differences and possibly pointing to yet unknown interactions between the transcriptomic regulation processes controlling the levels of these transcripts. Other studies have recently identified these 3 genes as regulators of neuronal morphology which modulate dendritic, axonal or synaptic changes. Also, entorhinal cortical thinning in Alzheimer’s disease patients has recently been attributed to a single nucleotide polymorphism (SNP rs11129640) residing in close proximity upstream to the Arpp21 gene^55^. While further investigation is needed to better link between the observed transcriptomic and morphological changes and disclose their cortical function, our current findings shed new light on the link between neuronal transcription and morphology and how they relate to each other.

### Cholinergic implications for age and disease states

Cholinergic networks are notably subject to differences between sexes and changes in mental disease states^27^, but the corresponding structure-function relationships are still unknown. Additionally, external stressors and inflammation inducers harm neuronal health and alter dendritic and synaptic structures^11^, again with unknown causes and characteristics. Perturbation of the cholinergic transmission modifies dendritic and synaptic morphology of cortical pyramidal neurons during late development^56^. Furthermore, large scale studies demonstrate elevated risk of dementia under prolonged exposure of aged individuals to anti-cholinergic medications^57^, and the cortical cholinergic innervation is severely impaired during the course of Alzheimer’s disease^58^ and under viral infection^11^. However, the corresponding changes in either the morphology or the transcriptomic profiles of cortical cholinergic neurons remain unknown. Combined multi-omic approaches such as those shown in our current study may shed light on these observations and enable the linking of specific neuronal subtypes and genes to elements of human health and wellbeing.

## Conclusions

We have developed a strategy for studying the baseline atlas and insult-induced plasticity in dendrite morphology of defined cortical VChI populations and disclose their morphological and transcriptomic resources. Recent advancements in the field of whole brain clearing, light-sheet imaging and single cell automatic reconstruction algorithms may expand this approach to fit denser neuronal populations and fine compartments such as dendritic spines and axons. We hope that our work can serve as a basis for progressing in the analysis and statistical approaches to achieve morphological profiling of reconstructed neuronal populations and combine these characteristics to the outcome of spatial transcriptomics^59^, progressing yet one more step ahead in high-throughput single cell resolution methods.

## Materials and Methods

### Mice

B57/B6 progeny mice derived from a cross of a loxP-Stop-loxP-tdTomato (Ai14 - Stock No. 007914, Jackson Laboratories) with ChAT-IRES-Cre mice (Stock No. 018957, Jackson Laboratories) were employed in this study.

### Ethics statement

All experiments were approved by the institutional animal care and use committees of The Hebrew University of Jerusalem (Ethics approval - NS-18-15456-3) which follow the National Research Council (US) Guide for Care and Use of Laboratory Animals. All experimental protocols were approved by the University Ethics Committee for Maintenance and Experimentation on Laboratory Animals, The Hebrew University, Jerusalem, Israel.

### Whisker Deprivation

For the whisker deprivation experiments, ChAT-CreXAi14 p7 pups were lightly anesthetized with isoflurane (1-2% by volume in O2, LEI Medical) for one minute then quickly removed from the anesthesia chamber and the entire whisker pad plucked by tweezers, or slightly pinched without plucking in control mice. After several minutes, once recuperated, treated pups were returned to the home cage and monitored until carried back to the parent cage. For labeling, on p14 mice were ear punched on the deprived side. For more information see supplementary table 4.

### Perfusion and Brain Dissection for iDISCO Clearing and Immunostaining

At p30 ChAT-CreXAi14 mice, both naïve and whisker deprived, were anaesthetized with isoflurane as above and were administered an intra-peritoneal injection of 200 mg/kg sodium pentobarbital. Following trans-cardiac perfusion with 1xPBS (5ml/min) and then 10% formaldehyde in 1xPBS (5ml/min), mouse brains were collected and further incubated O.N. at 4°C in 10% formaldehyde and then washed 3 times with 1xPBS for 30 min. The cortices were dissected and used for iDISCO+ clearing as described initially^15^ and further updated in https://idisco.info/idisco-protocol/. Briefly, during the iDISCO+ protocol, we stained the tdTomato expressing cells with an anti-RFP antibody (1:1200 dilution, Rockland, 600-401-379) for 5 days followed by Alexa-647 conjugated Donkey anti-Rabbit secondary antibody (1:1000 dilution, Jackson ImmunoResearch, 711-605-152) for 5 days.

### Microscopy

A light sheet microscope (UltraMicroscope II, LaVision BioTec) operated by the Inspector Pro software (LaVision BioTec) with fixed lens configuration 4x lens with 2x magnification was used. Images were acquired with a Neo sCMOS camera (2,560 × 2,160, pixel size 6.5 µm x 6.5 µm, Andor) in 16bit. Cortical samples were attached to the sample holder with epoxy glue and placed in a 100% quartz imaging chamber (LaVision BioTec). For whisker deprived mice, the light sheet was generated by a SuperK Super-continuum white light laser (emission 460-800 nm, NKT photonics) for auto-fluorescence (Ex 470nm/40nm) and VChIs (Ex 640nm/30nm). For naïve mice, the light sheet was generated by independent lasers (Module Gen. II, LaVision BioTec) for auto-fluorescence (488nm laser) and VChIs (639 nm). All autofluorescence scans were taken at estimated sheets of 10µm and 4µm z-step size with no horizontal focusing and signals were detected with Em 525nm/50nm filters. VChIs scans were taken at estimated 3.86µm sheet width and 1µm z-step size with 8 step horizontal focusing. Signals were detected with Em 690nm/50nm filters.

### Pop-Rec Pipeline

Each light-sheet scan was cropped such that only the cortex region was kept, and the barrel cortex region was traced and defined as an ROI (FIJI) based on normalized (Intensify3D) auto-fluorescent scans. Processed signal scans were normalized across experimental groups with (Intensify3D+) and neuron somata were detected using our algorithm (MATLAB) as follows. A cube (naïve - sized 350*350*500 or whisker deprived - 400*400*400 pixels) was isolated around each neuron and saved as individual 3D tiff files (MATLAB). Image cubes were fed to a MATLAB script running on a Linux cluster. Morphological features from candidate reconstructions were extracted for each cell (Python) and were filtered using Treender (MATLAB and Vaa3D^60^). Cortical surface and bottom border (Corpus Callosum) were measured from auto-fluorescent scans. All MATLAB and python codes are available in Codes.zip file.

### Morphology Features

Accurate definition of features described in this manuscript can be found at: https://neurom.readthedocs.io/en/stable/definitions.html

### Movies

Movie 1 and Movie 2 were edited and rendered with IMARIS (Bitplane). Movie 2 was initially filtered to remove bright noise puncta. Movie 3 was produced in FIJI.

### Statistical Analysis of morphological differences and clustering

All statistical analysis was performed with MATLAB (MathWorks). tSNE clustering was performed on 30 standardized (z-score, MATLAB) morphological features. To account for known biological variability such as repeated measures (animals) or animal sex we used a linear mixed effects approach as described in https://www.mathworks.com/help/stats/fitlme.html. Exact models are described in Supplementary Tables 3 and 4.

### Single Cell Tissue Dissociation

To obtain samples for RNA-Seq, whisker deprived mice were sacrificed at day 30 by isoflurane anesthetizing (as described above) followed by cervical dislocation. The left and right barrel cortices were dissected and subjected to the NuNex procedure (Yayon et al., manuscript in preparation). Briefly, tissues were dissected in ice-cold HBSS (Sigma-Aldrich H6648), immediately transferred to 1ml of 10% formalin solution (Sigma-Aldrich HT501128) and incubated at R.T. for 30 minutes. Samples were then washed in 1xPBS, chopped with a scalpel and homogenized with a Dounce tissue grinder (Sigma-Aldrich, D9063). The homogenate was passed through a 40µm cell strainer (BD Falcon) and pelleted by centrifugation at 900 x g for 5 min. Cells were resuspended and stained with DAPI (Santa Cruz) for 10min. All steps were performed on ice.

### VChI FACS Enrichment

To enrich for barrel cortex VChIs, we sorted cells based on DAPI and tdTomato signals. We combined cells from several mice to reach roughly 200 VChIs per pool, ensuring each pool was of the same cortical side, including the same mice, treatment condition and gender. To ensure reproducible experimental conditions we limited whisker plucking to the right side only. RNA was extracted from the pools of sorted cell with a dedicated kit for formalin-fixed tissues (Qiagen, 217504) according to manufacturer’s instructions. See supplementary Table 5 for details on sample characteristics and RIN values.

### Library preparation and Sequencing

800pg total RNA from each sample pool was subjected to library construction according to manufacturer’s instructions (SMARTer® Stranded Total RNA-Seq Kit v2 - Pico Input Mammalian, Takara Bio). Libraries were barcoded, pooled and sequenced with a NextSeq 550 Sequencer (Illumina).

### RNA-Seq Analysis

The analysis pipeline included adapter trimming with cutadapt (https://cutadapt.readthedocs.io/en/stable/) followed by alignment to the mouse genome with STAR61 and detection of differential expressed genes with DESeq262 under R studio (https://rstudio.com/).

### Single Cell Data

Data was downloaded and analyzed from (https://github.com/AllenInstitute/scrattch.hicat) with custom R packages. Counts from VChIs were normalized to counts per million and subjected to tSNE clustering on standardized z-score data (MATLAB) of the top 5000 expressed transcripts.

## General

The authors are grateful to Sebastian Lobentanzer, Frankfurt, and Shani Vaknine, Jerusalem for advice and assitance in RNA-Seq analysis; and to Drs Yoseph Addadi, Rehovot, and Ester R Bennett, David S Greenberg, Sagiv Shifman and Ron Refaeli, Jerusalem for their contributions towards this study.

## Author Contribution

**N**.**Y**. performed entirely or participated in all experimental procedures and analyses including MATLAB, FIJI and R coding. **O**.**A**. and **N**.**Y**. wrote the code for cluster-based reconstructions and **O**.**A**. wrote the codes for feature extractions (Python, NeuroM) and reconstruction sorting and provided computational advice and coding throughout the progression of the project and writing of the manuscript. **A**.**D**. assisted in animal procedures, VChIs morphology estimation and writing of the manuscript. **O**.**Y**. Performed the procedures for VChI RNA extraction including homogenization steps and FACS sorting. **G**.**A**. Produced a Rhino pipeline for 3D large scale rendering of VChIs for visualization purposes and advised on the Intensify3D+ algorithm. **M**.**L**. and **I**.**S**. guided **O**.**A**. and **A**.**D**.’s contributions. **M**.**T**.**L** and **N**.**R**. Initiated the project, guided **N**.**Y**. in strategic planning of iDISCO clearing and Light-Sheet imaging experiments. **N**.**R**. also participated in preparation of the manuscript. **H**.**S**. oversaw and guided the project, preparation and writing of the manuscript.

## Data and materials availability

upon request and/or prior to publication all codes and data used to generate the figures will be publicly available by a GitHub repository. Due to size, original light-sheet scans (TB is size) will be sent upon specific request. Due to size, original fastq files used for RNA-Seq analysis will be sent upon specific request.

