## Supplemental Figures and Movies for "Morphometric reconstructions atlas shows insult-driven plasticity in cortical VIP/ChAT interneurons"

**­­­­­­­**

­**Supplementary Figures and Movies**

**Movie 1.** Whole iDISCO+ cleared mouse brain showing labeling of Cholinergic neurons and projections.

**Movie 2.** Representative example of a 400µm-thick section and PopRec reconstructions of 207 VChIs presented in random color for clarity.

**Movie 3.** Representative example of a 4x2mm-thick section and macro-feature extraction: the location of the surface of the cortex (grey) and barrel fields (green). In addition, the rendering of large blood vessels is represented (red). All features were extracted from autofluorescence without additional staining.

Figure S1. Graphical user interface (GUI) Manual for intensify3D+

For the full code, tutorials and theory please see:

<https://github.com/nadavyayon/Intensify3D>

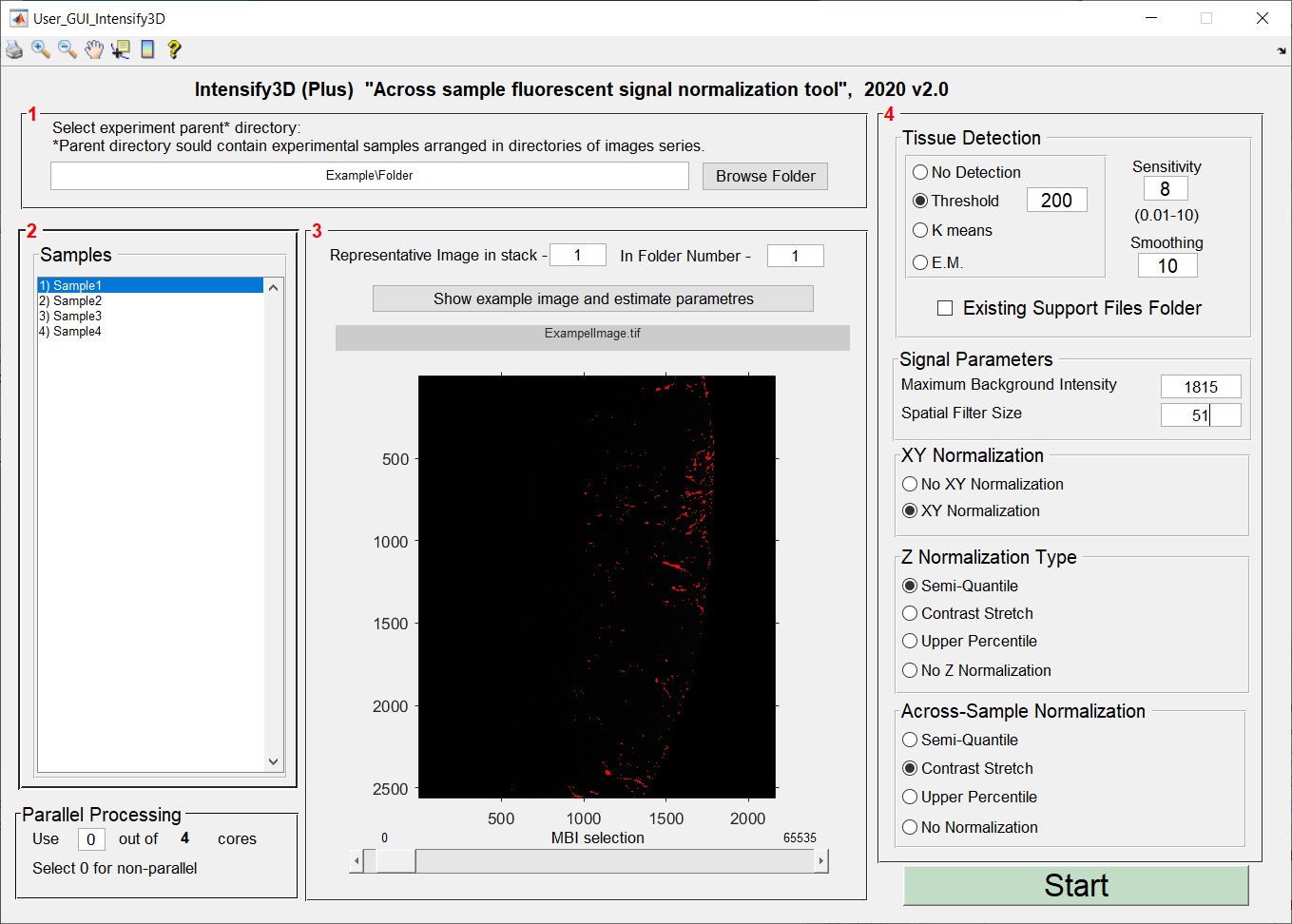

**General Recommendations**

Before starting please read the Intensify3D original manuscript and make sure the assumptions of normalization are met. Intensify3D+ can correct an unlimited number of images from an unlimited number of samples since it operates in a serial manner. Hence, it only supports image sequences. . The *.tiff files should ideally be unprocessed data in a 12 or 16bit format. Memory requirements depend on image size and parallel processing. Based on our experience, the maximum requirements are 750 bytes/pixel. For example, processing a single Light-Sheet image of 2560x2160 pixels will require ~ 4Gb of RAM from each processor + 4Gb for general processes. For example, if your PC has 4 cores, it is possible to analyze 4 Light Sheet images simultaneously, which will require 20Gb of RAM. It is highly recommended to start with a few representative images (~20), adjust the parameters and only then run the process on the entire stack or samples.

**Operation instructions and GUI options:**
The graphical user interface is divided to 4 panels:

**Panel 1** – Parent folder selection: Here the user selects the directory containing the different biological sample set divided into directories.

**Panel 2** - Shows which directories were found and will be normalized withing and across samples.

**Panel 3** – Estimate your background: The objective of this section is to assist the user in selecting the ideal maximum background intensity (MBI) in a single image. This value will be used by Intensify3D+ to estimate the background across all images in the stack. “Image number” and “Folder Number” are used to select a representative image from the stack that carries a clear signal. Once the image has been selected, pressing the “show image and estimate parameters” button displays the requested image is displayed, a brightness contrast adjustment window opens and an initial estimation of the MBI is assigned based on the 100th percentile of intensity potentially showing only signal pixels in red. Next, the user should adjust the MBI selection with the dedicated slide bar at the bottom of the image in the following order: (1) adjust brightness and contrast (2) move slide bar to set MBI (performing 2 before 1 will show a distorted selection of the MBI). Repeat 1->2 until satisfied with the result*. The matched value for the MBI will be set in the “stack parameters” section in panel 4.

**Panel 4** – Setting run parameters:

Automatic Tissue Detection - Intensify3D+ has the ability to detect the background or tissue area in an image in 2 ways: simply by taking all the pixels above a certain threshold as the tissue, or by clustering algorithms: K means and Expectation Maximization (E.M.). This option is critical for images where not all the image area is relevant for normalization. The sensitivity of the tissue detection should be tested by the user to fit to the specific image set.

Parallel processing section: Is very useful when analyzing large image stacks: the GUI detects how many cores your CPU has and offers the user the option of how many of them will be dedicated to the run. If your MATLAB license does not include the parallelization package, select 0 and work without it (this limitation does not apply to the standalone version).

Signal parameters section: Select the MBI* (described below) and the spatial filter size which determines the spatial frequency of gradient background that should be corrected. Spatial filter size determines the frequency of background gradients to be corrected by Intensify3D+. The minimum value for this parameter should be a least twice the diameter of the largest signal structure. Lower values could affect the signal.

XY Normalization: Determine whether to normalize the XY direction of each image.

Z normalization type section: Allows section of the desired normalization type across the images in the stack. Semi-quantile is the “strongest” correction converting the pixel histogram to match across the entire stack. Contrast-Stretch matches 2 values (lower and upper quantiles) of the intensity histograms across the stack. Upper percentile matches a single value across the stack and No Z Normalization does not correct between images of the stack. (for more information see Intensify3D manuscript). Last, (for more information see main text and supplementary figure S4).

**Last, after running Intensify3D+, follow the command window in MATLAB to estimate progress.**

*intuition for selection of the MBI value: The correct approximation of the image background depends on “cleaning out” the signal pixels by thresholding and spatial filtering. High brightness signal pixels can affect the ability of the spatial filter. The MBI should be set so that the most signal pixels will be removed without removing background pixels. Notice the red-labeled pixels in the example image of bright spheres. Lowering the MBI would result in removing background pixels and increasing the MBI would retain more signal pixels. Both ways will lead to a sub-optimal estimation of the background

**Supplementary Figure S2. Tree-nder ranking and quality control algorithm.**

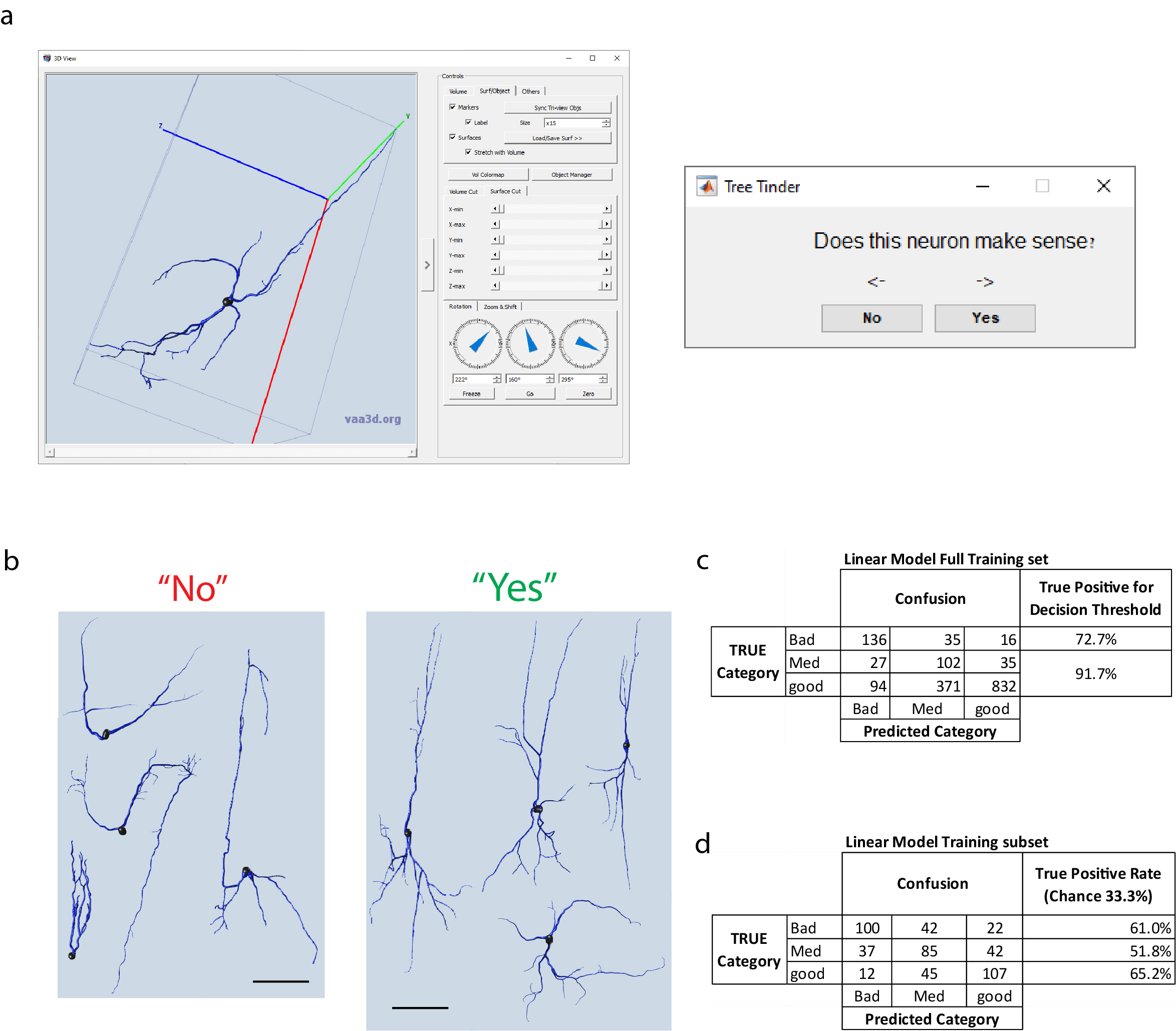

**a**. Graphical user interface for Treender quality control algorithm. Left – Vaa3D window shows the candidate neuron for estimation in 3D. Right – MATLAB interface that is automatically displayed prompts for selection using right and left keystrokes. **b**. Example neurons that were classed as “bad” and “good” according to their general structure. **c**. Accuracy of regression linear model on all neurons in training set and based on quality threshold. Cells that are predicted to be “Bad” are excluded. **d.** Accuracy of regression linear model on balanced training set.
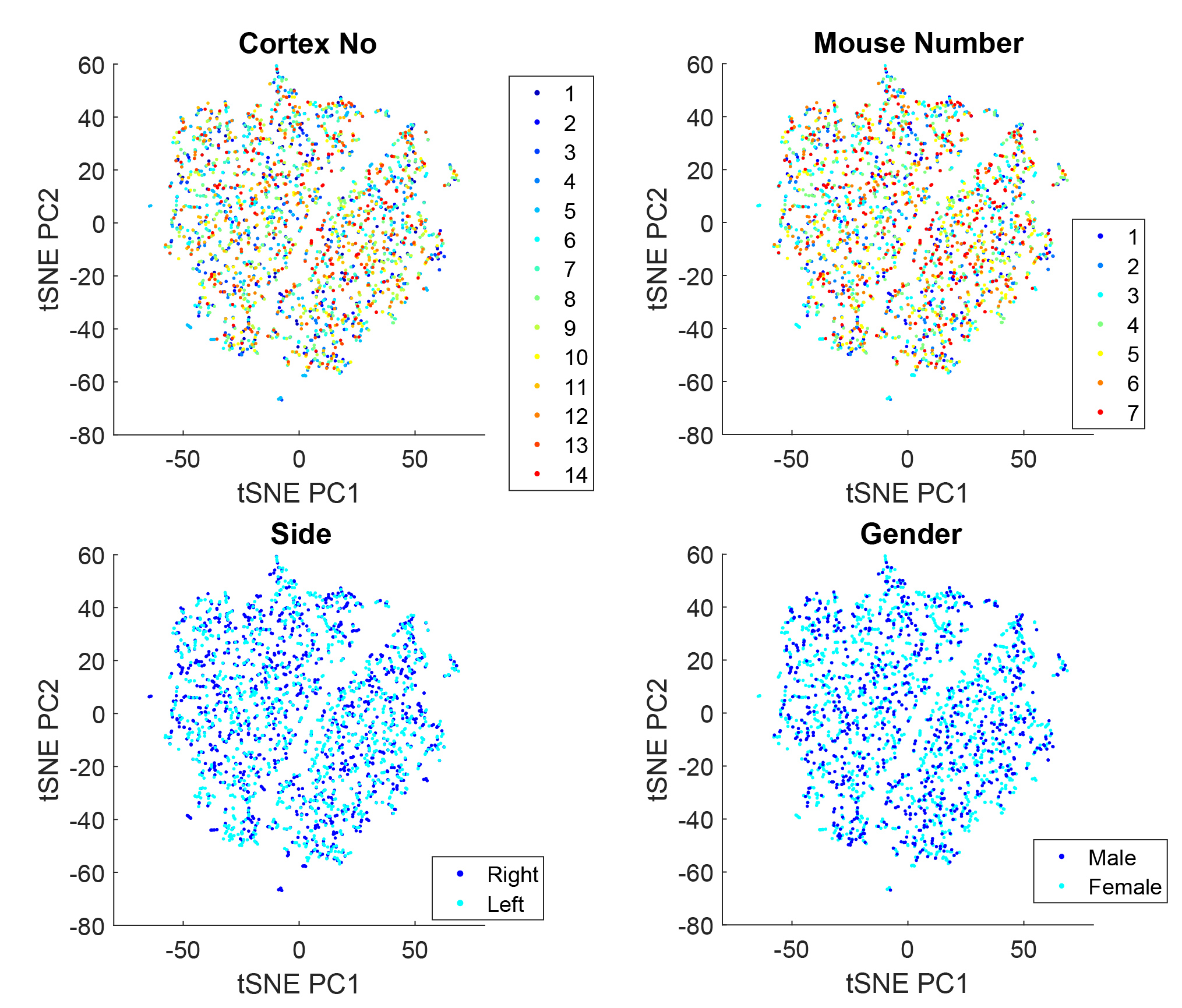
**Supplementary Figure s3. TSNE on metadata**

tSNE analysis of VChIs with cells tagged according to 1) cortex number (sample number), 2) mouse number, 3) cortical side or 4) animal sex. For all these parameters, we saw no clear sub-clustering, indicating that the original clustering is maintained also when considering experimental, sex and hemisphere variability.

Supplementary Figure 4

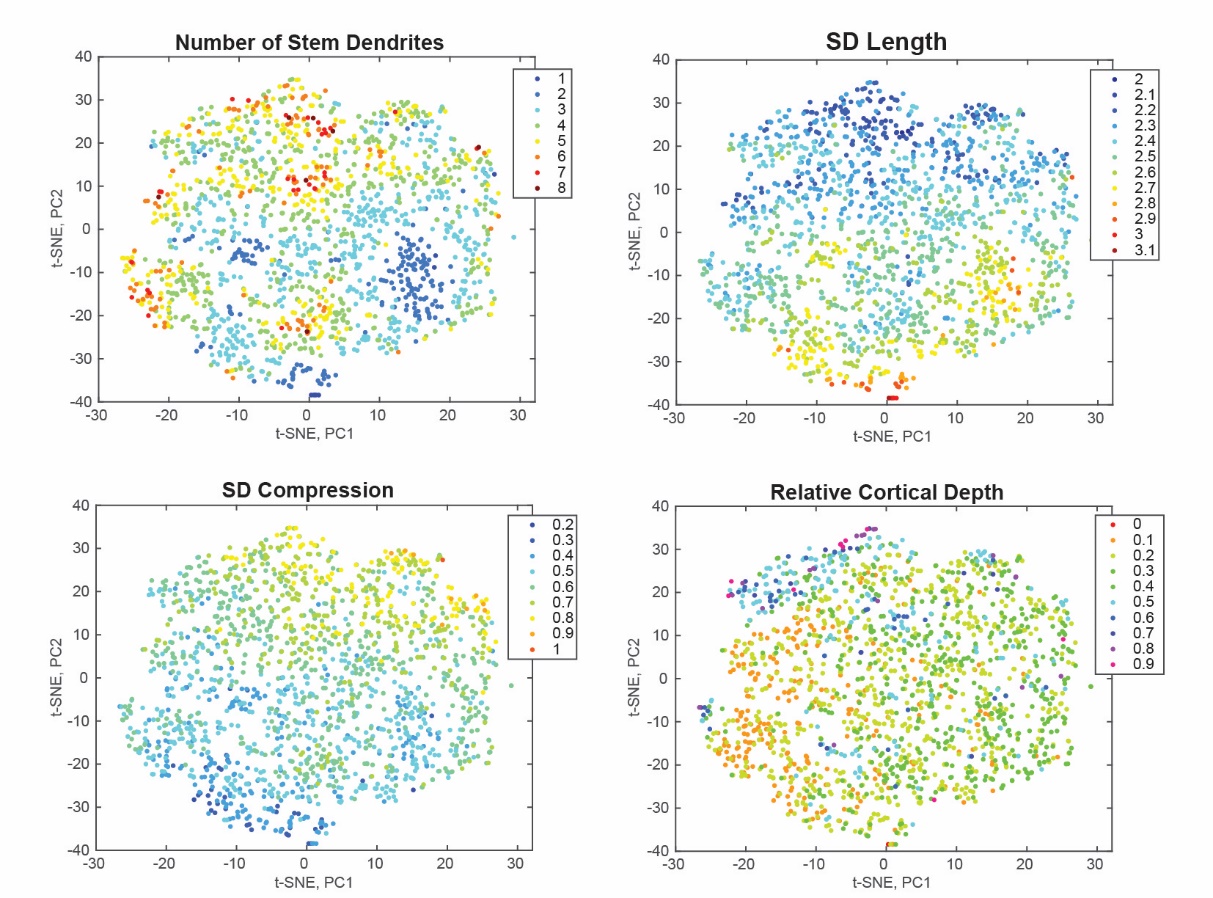

Supplementary Figure 4. tSNE clustering of whisker deprivation batch shows no clear clustering according to number of stem dendrites. This sample set showed a significant halo around the cell soma which might have interfered with accurate detection of stem dendrites (see materials and methods, Microscopy). Alternatively, this could be a result of experimental manipulation. For these reasons we focused on quantification of changes in the entire VChI population.
